# Satellite DNA Editing Enables Meiosis-Independent Chromosome Engineering

**DOI:** 10.64898/2026.06.02.729711

**Authors:** Ran Zhou, Margot S.S. Chen, MaKenzie R. Drowns, Kathrine Mailloux, Kangquan Yin, Brieanne Vaillancourt, C. Robin Buell, Chung-Jui Tsai

## Abstract

Chromosome□scale genome engineering in plants typically relies on locus-specific recombination during meiosis to select desirable outcomes, limiting its application in species with long generation times. Here, we show that CRISPR targeting of abundant satellite DNA enables multiplexed chromosome restructuring in aspen, generating diverse large-scale structural variants in the first generation. These megabase rearrangements remain mitotically stable through clonal propagation and regeneration with no obvious growth defects.

## Main text

Chromosome engineering offers a route to reshape genome architecture and alter recombination, gene dosage, or karyotype, with clear relevance to agriculture [1]. To date, however, chromosome□scale engineering has been largely confined to model plants, where targeted double□strand breaks (DSBs) enable defined intra- or interchromosomal rearrangements, typically followed by multiple rounds of genetic crosses to select desired outcomes [2–8]. In long-lived tree species, with extended juvenile phases and limited opportunities for crossing, this strategy remains impractical.

Here, we demonstrate a chromosome engineering strategy in *Populus tremula × alba* (INRA 717-1B4, hereafter 717) by leveraging CRISPR–Cas9 for hi-plex, multi-locus editing of M147, an abundant, aspen□specific satellite DNA. M147 is present in non□centromeric tandem repeat arrays (TRAs) on most chromosomes in 717 [9], such that a single guide-RNA (gRNA) targeting the consensus sequence can induce simultaneous DSBs at many loci across both haplotypes. Combined with tissue culture–regeneration, this enables chromosome-scale engineering and selection in a perennial tree, bypassing genetic crosses (Figure 1A). We tested this approach using two gRNA designs, M147.g7 and M147.g8, with 47,965 and 29,853 predicted target loci in the 717 genome, respectively (Table S1). Stable transformation by *Agrobacterium tumefaciens* with these constructs yielded 14 and 27 independent callus lines, of which, 2 and 3, respectively, regenerated into whole plants—well below typical recovery rates [10]. Callus recovery was lower for M147.g7 with a higher number of target sites, suggesting reduced tolerance to extensive genome cleavage. A second transformation with M147.g8 using triple the number of leaf-disc explants yielded 50 calli and 11 independent plant lines (Table S2), demonstrating reproducible, albeit low, transformation efficiency.

**Figure 1.**
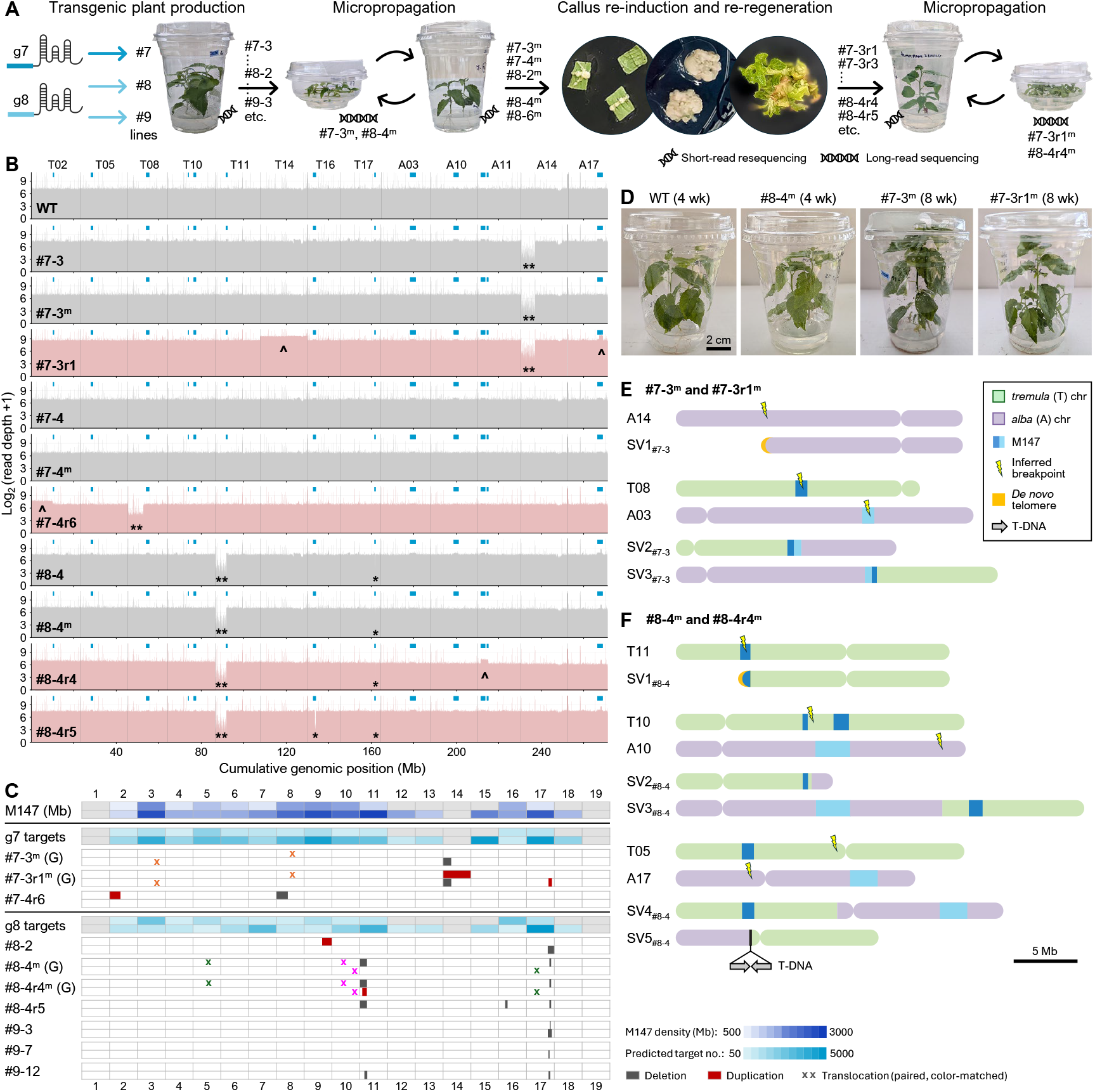
Genome-wide structural variation in transgenic *Populus*. **A**, Overview of transgenic lines and sampling scheme. **B**, Resequencing read coverage (log□) for affected chromosomes in WT and transgenic lines (m, micropropagated; r, re-regenerated). Blue bars indicate M147-TRAs (*, deletion; **, arm-loss; ^, duplication). **C**, Summary of large-scale structural variants across 38 chromosomes (*tremula*, top; *alba*, bottom) inferred from short-read mapping or long-read genome assemblies (G). Heatmaps show M147 abundance (dark blue) and target density (light blue); deletion (black), duplication (red), translocation (paired ×). **D**, Representative plants in tissue culture. **E–F**, Structural variants revealed by long-read sequencing (SVs labeled). Green, *tremula*; purple, *alba*; blue, M147-TRAs; yellow, *de novo* telomere; lightning, inferred breakpoint; arrow, T-DNA insertion. Scale bar, 5 Mb.

Whole-genome resequencing using short reads revealed discernible structural variants (SVs) in six of the 16 regenerated lines (37.5%), with the remaining lines indistinguishable from wild type (WT). These SVs included deletions, chromosomal arm-loss, and a segmental duplication, corresponding to 10 loci ranging from 130 Kb to 6.7 Mb (Figure 1B–C, Table S3). Four lines showed two distinct SVs occurring either on homologous *P. tremula* (T) and *P. alba* (A) chromosomes (line #9-3) or on non-homologous chromosomes (#8-2, #8-4, #9-12, Figure 1C). In most cases, the affected intervals overlapped M147 TRAs (blue ticks, Figure 1B), with a slight preference for chromosomes carrying the highest densities of predicted M147.g8 target sites, including A11 and A17 (Figure 1C). This pattern is consistent with CRISPR-induced SVs at regions of high target density. A notable exception was #7-3, which harbored a 6.7 Mb arm-loss on A14—a chromosome devoid of M147 [9] (Figure 1B–C). Given that M147.g7 is predicted to impose a massive genome-wide cleavage burden, this event may reflect collateral chromosomal instability arising from extensive DNA damage rather than direct targeted cleavage.

To expand structural diversity without crossing, five lines that had been micropropagated (*m*) were subjected to callus re-induction followed by shoot regeneration (Figure 1A). We recovered 22 re-regenerated (*r*) plants (2–7 per line, Table S2). Under tissue culture conditions, plant growth appeared similar among WT and transgenic lines, regardless of culture history (Figure 1D). Whole□genome resequencing of the micropropagated lines showed stable parental read-mapping patterns with no additional large-scale SVs (Figure 1B). In contrast, new SVs were detected in four re-regenerated plants (18%) at six loci (Table S3), with the remainder indistinguishable from their parental lines. These data suggest dedifferentiation into mitotically active callus facilitated further Cas9–mediated genome restructuring.

The new SVs from re-regenerants included M147-TRA deletions and duplications, arm-loss, arm-duplication, and a whole-chromosome duplication (Figure 1B–C, Table S3). As in parental lines, most affected loci either overlapped or flanked M147-TRAs, with estimated sizes ranging from 590 Kb to 9.9 Mb. The exception was the #7-3r1 whole-chromosome duplication of T14, the homologous chromosome of A14 which had undergone arm-loss in the parental line. Because both A14 and T14 lack M147 gRNA target sites, this whole-chromosome duplication likely represents a compensatory response during callus re-induction to restore gene dosage lost in the parental line.

To more thoroughly assess genome engineering outcomes, we performed long□read nanopore sequencing of two line-pairs (#7-3 and #8-4), along with WT-717. Each pair comprised parental and re-regenerated lines, both derived from micropropagated material (Figure 1A), yielding five *de novo*, haplotype-resolved, telomere-to-telomere genome assemblies (Table S4). Parental SVs identified using short-read resequencing were confirmed and breakpoints resolved, including a 6.7 Mb arm-loss on A14, a 5.3 Mb arm-loss on T11, and a 380 Kb deletion within an M147-TRA on T17 (Figure 1C). Both arm-loss events were associated with *de novo* telomere formation, evidenced by canonical telomere repeats immediately distal to the inferred breakpoints. Estimated telomere lengths reached ~28 Kb on truncated A14 (SV1_#7-3_) and ~17 Kb on truncated T11 (SV1_#8-4_, Figure 1E–F, Table S5).

Three SVs uniquely associated with callus re-induction were also confirmed, including two M147 duplications (1.5 Mb on A17 in #7-3r1 and 3.4 Mb on A11 in #8-4r4) and a T14 whole-chromosome duplication in #7-3r1 (Figure 1C). The structural configuration of the duplicated T14—whether present as a free chromosome or fused to another chromosome—could not be resolved.

Long-read assemblies further uncovered cryptic rearrangements not detected by reference-based short-read mapping. Both #7-3 and #7-3r1 harbored an inter-chromosomal translocation between T08 and A03, with breakpoints mapped within M147-TRAs, consistent with Cas9–mediated cleavage. This translocation produced two chimeric chromosomes (Figure 1E): a ~17 Mb derivative (SV2_#7-3_) carrying the T08 centromere (CEN-T08) and a ~25 Mb derivative carrying CEN-A03 (SV3_#7-3_).

Both #8-4 and #8-4r4 assemblies revealed two cryptic rearrangements. The first involved homologous T10 and A10 with non-syntenic breakpoints: the T10 breakpoint occurred between two small M147-TRAs, while the A10 breakpoint mapped to the distal long-arm (Figure 1F). This translocation generated a 12.6 Mb CEN-T10 chromosome (SV2_#8-4_), the smallest in the genome, and a 32 Mb CEN-A10 chromosome (SV3_#8-4_), the third largest. The second translocation, between T05 and A17, involved pericentromeric breakpoints (Figure 1F) and produced a ~25.7 Mb chromosome carrying two M147 TRAs (SV4_#8-4_) and a 16.5 Mb M147-free chromosome (SV5_#8-4_). The 16.5 Mb derivative contained a head-to-head insertion of two T-DNA copies at the fusion junction (Figure 1F), suggesting that at least one breakpoint coincided with T-DNA integration. In all cases, shared rearrangements between parental and re□regenerated samples indicate mitotic stability during micropropagation and callus re□induction followed by organogenesis.

Together, these results demonstrate that CRISPR–Cas9 targeting of abundant satellite DNA can drive chromosome-scale restructuring in a woody perennial without reliance on meiotic crossing. This platform generated diverse structural outcomes even from limited sampling of transgenic lines, with individuals harboring up to seven distinct SVs. The extent of SVs is likely underestimated, as many rearrangements are cryptic to short□read approaches. Importantly, these SVs remain mitotically stable through micropropagation and callus re□regeneration. Our findings extend chromosome engineering beyond annual crops and establish satellite DNA as a tractable target for probing genome structural plasticity at scale. Given the widespread occurrence of high-copy satellite DNA in eukaryotic genomes, this strategy should be broadly applicable across species. Finally, this work provides a germplasm resource for dissecting chromosome dosage effects and exploring genome minimization in long□lived, clonally propagated tree species.

## Declarations

### Ethics approval and consent to participate

Not applicable

### Consent for publication

Not applicable

### Data availability

Illumina short-read and Nanopore long-read sequencing data have been deposited in the NCBI Sequence Read Archive under BioProject PRJNA1469888.

### Authors’ contributions

CJT and RZ designed the research, MSSC and MRD performed plant experiments, KM, KY, and RZ conducted sequencing. RZ performed data analysis with assistance from BV and CRB. RZ and CJT wrote the manuscript with input from all authors.

## Acknowledgements

We thank Magdy Alabady and McKenna Coile for their assistance with genome resequencing and the Georgia Genomics & Bioinformatics Core (RRID:SCR_010994) for Illumina sequencing.

